# Optimal distributions of receptors on arbitrarily shaped cell surfaces

**DOI:** 10.1101/2025.03.15.640757

**Authors:** Daoning Wu, Sayun Mao, Jie Lin

**Author notes:** These authors contributed equally to this work (Daoning Wu, Sayun Mao).

## Abstract

Efficient absorption of signaling molecules and carbohydrates through receptors on cell surfaces is crucial for various biological processes. While ubiquitous patterns of receptor distributions, including polar localization in rod-shaped cells, have been widely observed experimentally, their underlying evolutionary advantage is unclear. In this work, we study how spatial distributions of receptors on cell surfaces affect the total flux entering the cell. We innovate a method by which one can calculate the fluxes through all receptors using linear equations, which applies to arbitrarily shaped cells. Our theories recover previous results for spherical cells and further show that the flux through each receptor is spatially dependent in non-spherical cells. In particular, the fluxes are the highest near the poles in rod-shaped cells and the highest near the invagination in defective spherical cells. Surprisingly, we prove that the optimal receptor distribution on an arbitrarily shaped cell maximizing the total flux is precisely the charge density distribution on an ideal conductor of the same shape, which agrees with numerical simulations. Our work unveils the evolutionary origin of receptor localizations.

---

Many intracellular processes rely on the arrival of essential molecules at the cell surface, where the cells absorb the molecules via specific membrane receptors [1, 2]. For example, cellular metabolism requires the cell to uptake carbohydrates efficiently [3, 4], e.g., through the phosphotransferase system (PTS) in bacteria [5–7]. For rod-shaped bacteria, e.g., *Escherichia coli*, experiments found that many types of receptors (or transporters) are localized at the cell poles, e.g., the osmosensory transporter ProP [8], the lactose transporter LacY [9], the PTS proteins EI and HPr [10], and the chemotaxis receptors [11, 12]. Interestingly, in defective spherical *E. coli* cells with an invagination, the PTS proteins are instead localized at the edge of the invagination [13]. Mechanically, polar constituents, e.g., anionic lipids [8, 14], and curvature sensor proteins [15] may serve as geometric cues for polar localization. In this work, we seek to understand the localizations of receptors from an evolutionary perspective: what kinds of spatial distribution of receptors maximize the total molecule flux entering the cell given a fixed environment? We propose that achieving the maximum absorbing flux may be the evolutionary driving force for certain distributions of receptors ubiquitous in biology.

The upper bound of flux is achieved when the entire cell surface is a perfect absorbing boundary so that any molecule hitting the surface is absorbed. For a spherical cell, the upper bound of molecule flux *J*_max_ = 4*πDRc*_∞_where *D* is the diffusion constant of the molecule of interest, *R* is the cell radius, and *c*_∞_ is the molecule concentration far from the cell. Phillips et al. modeled a spherical cell surface as a homogeneous non-absorbing boundary [16]. The total flux *J* = *Nk*_on_*c*(*a*) where *N* is the number of the receptor, *k*_on_ is the absorption rate of each receptor, and *c*(*a*) is the molecule concentration on the cell surface, which is determined self-consistently. They showed that the molecule flux entering the cell is a Hill function of the receptor number: 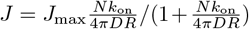,consistent with the experimental measurements of the adsorption rate of bacteriophage *λ* on the number of *λ*-receptors in *Escherichia coli* [17]. By taking *k*_on_ ≈ 10 *µ*M^−1^s^−1^, they showed that the receptors only need to cover about 0.1% of the cell surface to achieve a flux 50% of the upper bound. This surprising finding leads to the conclusion that cells only need to allocate a small fraction of their surfaces to one receptor type to achieve a high absorption efficiency, allowing them to simultaneously express multiple types of receptors.

The previous analyses by Phillips et al. were inspiring. Nevertheless, critical questions remain. First, the actual boundary condition on the cell surface is complex since receptors only cover a finite fraction. Most importantly, the effects of non-spherical cell shape on the molecule flux are far from clear. For an arbitrarily shaped cell, what is the optimal distribution of receptors maximizing the flux entering the cell? In this work, we answer these questions by studying an arbitrarily shaped cell with its surface decorated by multiple receptors. Each receptor is modeled as a circular patch with an absorbing boundary condition. Berg and Purcell have studied a particular case of uniformly distributed receptors on a spherical cell [18]. Our key theoretical innovation is to convert the problem of solving Poisson’s equation with complex boundary conditions to a linear algebra problem involving the flux of each receptor, applicable to arbitrarily shaped cells with nonuniformly distributed receptors. This method allows us to compute the fluxes of all receptors exceedingly faster than other numerical methods, e.g., finite element simulations [19]. For spherical cells, we recover Berg and Purcell’s classical results [18], which strongly supports the validity of our method; in particular, we relate the absorption rate *k*_on_ introduced by Phillips et al. to the receptor size, *k*_on_ = 4*Dr* where *r* is the receptor radius and explicitly demonstrate that only a tiny fraction of area is needed to achieve a molecule flux close to the upper bound.

For an arbitrarily shaped cell with uniformly distributed receptors, the fluxes of receptors depend on their locations. For a rod-shaped cell, the fluxes of receptors at the poles are higher than those on the side. For a spherical cell with a bud, the fluxes are the highest at the bud tip. Surprisingly, for a spherical cell with an invagination, the fluxes at the edge of the invagination are the highest on the cell surface, rationalizing the localization of the PTS proteins observed in defective spherical *E. coli* [13].

For an arbitrarily shaped cell with nonuniformly distributed receptors, we prove that the optimal distribution of receptors that maximizes the total flux is precisely the charge density distribution on an ideal conductor of the same shape. We test our predictions by simulating an evolutionary process in which we let the locations of receptors evolve to maximize the total flux. As predicted, the receptor distribution converges to the charge density distribution on an ideal conductor. Our theories provide a physical foundation for the evolution of receptor localizations, e.g., rationalizing the ubiquitous polar locations of receptors in rod-shaped cells.

## Computing fluxes for an arbitrarily shaped cell

We study an arbitrarily shaped cell with *N* circular receptors decorating its surface Ω. The boundary condition of the concentration field is that *c*(***x***) = 0 for ***x*** ∈ *S*_*i*_ where *i* = 1, 2, …, *N* and *S*_*i*_ is the region occupied by receptor *i*. For *x* outside any receptors, 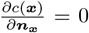.Here, ***n***_*x*_ is the unit vector perpendicular to the surface at ***x*** pointing to the cell interior. The flux through a receptor is 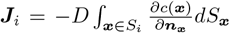 .Because the radius of typical receptors *r* ∼1 nm, the average distance between nearby receptors *l*∼ 10 nm [20], and the typical size of a bacterial cell *R* ∼1000 nm [21], we use the condition *r* ≪*l*≪ *R* in the following, a good approximation for most biological scenarios of our interest.

We use the implicit solution for Poisson’s equation [22] and adapt it to the case where the concentration at infinity is *c*_∞_

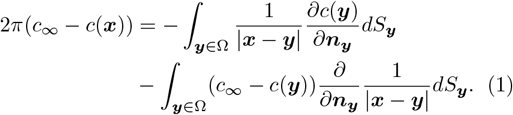

Here, 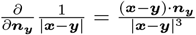. We take ***x*** =***y****i* in Eq. (1),

where ***y***_*i*_ is the center of receptor *i*, and rewrite Eq. (1) as

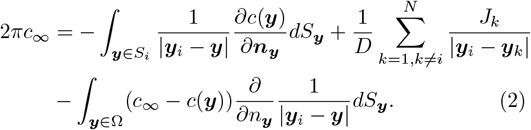

Here, we use the definition of flux to replace the integrals over all the receptors except *i* by their fluxes. The first term on the right side of Eq. (2) can be considered as the electric potential Ψ of a flat disk (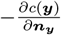can be considered as the charge density); therefore, it is proportional to the flux by the capacity 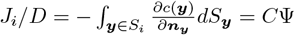 where *C* = 2*r/π* is capacity of a thin disk [23].

We next sample *M* auxiliary points uniformly on the cell surface that are not close to any receptors, which is always doable under our assumptions. We define *C*_*j*_ = *c*_∞_ − *c*(***x***_*j*_) where *j* = 1, 2, …*M* . Using Eq. (1), we find that

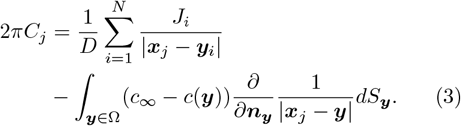

We approximate the integral over the cell surface in Eqs. (2, 3) as a sum over the *M* auxiliary points, i.e., 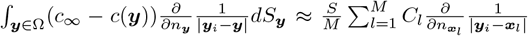 where *S* is the total surface area. Combing Eqs. (2, 3), we obtain the linear equations to solve the fluxes of all receptors:

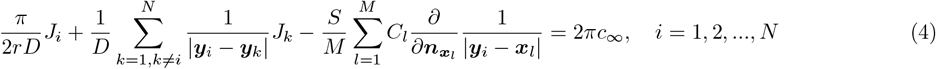

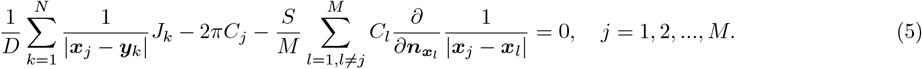

One should note that the *l* = *j* term in the last term of the left side of Eq. (5) is negligible in the continuous case; therefore, it is not included in the discrete summation.

## The spherical cell

We first study a spherical cell with its receptors uniformly distributed on its surface. In this

case, the flux through each receptor must be equal, and the concentration field at locations not close to any receptors must be uniform, that is, *J*_1_ = *J*_2_ = … = *J, C*_1_ = *C*_2_ = … = *C*. We approximate the summations 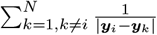 in Eq. (4) and 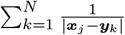 in Eq. (5) as 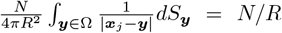. Similarly, we approximate the summations 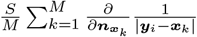 in Eq. (4) and 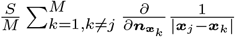 in Eq. (5) as 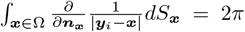. Given these simplifications due to the spherical shape, we obtain the total flux over all receptors as

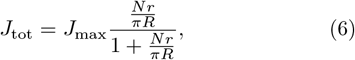

where *J*_max_ = 4*πDRc*_∞_, the total flux when the entire cell surface is absorbing boundary. Our results perfectly recover Berg and Purcell’s classical results for spherical cells [18], supporting the validity of our method.

By comparing Eq. (6) with the results of Phillips et al., we find the microscopic expression of the effective absorption rate *k*_on_ = 4*Dr*. Notably, the flux only depends on the total perimeter of all receptors, which suggests that the flux density is localized on the rim of each receptor, in agreement with the result of a circular absorbing trap on a flat surface [24]. We introduce *ϕ* = *Nr*^2^*/*4*R*^2^ as the fraction of total surface area occupied by receptors, and our results explicitly demonstrate that as long as *R/r* ≫ 1, the receptors only need to cover a small fraction of surface area for the total flux to be comparable to the upper bound.

We verify Eq. (6) by sampling *N* receptors uniformly on a spherical cell using the Fibonacci lattice method [25] (Figure 1a). In this work, we always uniformly sample *M* = *N* auxiliary points unless otherwise mentioned. We solve the linear equations Eqs. (4, 5). As expected, the fluxes are uniformly distributed, that is, the fluxes are the same across different receptors. The numerical total fluxes agree well with the theoretical predictions, and the total flux only depends on *Nr/πR* (Figure 1c). To test our methods’ robustness, we randomly sample *N* receptors on the cell with a uniform density: the average number per unit area is constant. To show the spatial dependence of fluxes, we plot the accumulated fluxes of multiple independent samplings (Figure 1b), which are also uniform. The total fluxes also agree well with the theoretical predictions in this random case (Figure 1c).

**FIG. 1.**
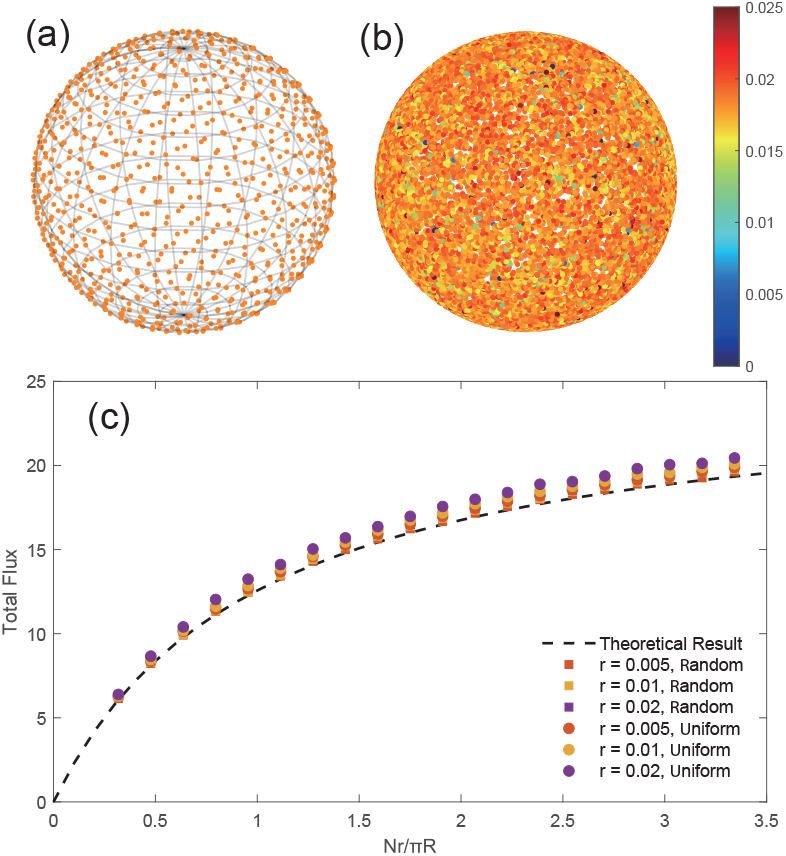
(a) We uniformly sample *N* = 800 receptors with radius *r* = 0.01 on a spherical cell with radius *R* = 2. (b) The accumulated fluxes of 20 independent samplings where the receptors are randomly sampled with a uniform density. (c) We change the number of receptors *N* for a given *r*, and the calculated total fluxes agree well with the theoretical predictions for both uniformly and randomly distributed receptors. In this work, the color of each receptor represents the flux through it.

## Arbitrarily shaped cells with uniformly distributed receptors

We next study the cases of nonspherical cells with uniformly distributed receptors. For rod-shaped cells, e.g., *E. coli*, we find that the fluxes are higher in the receptors near the poles than those on the side, and as the cell length increases, the difference becomes more significant (Figure 2). These observations explain the ubiquitousness of the polar localization of receptors in rod-shaped cells. Later, we show that if we allow receptors to move on the cell surface to maximize the total flux, they are enriched at the cell poles, which agrees with experimental observations.

**FIG. 2.**
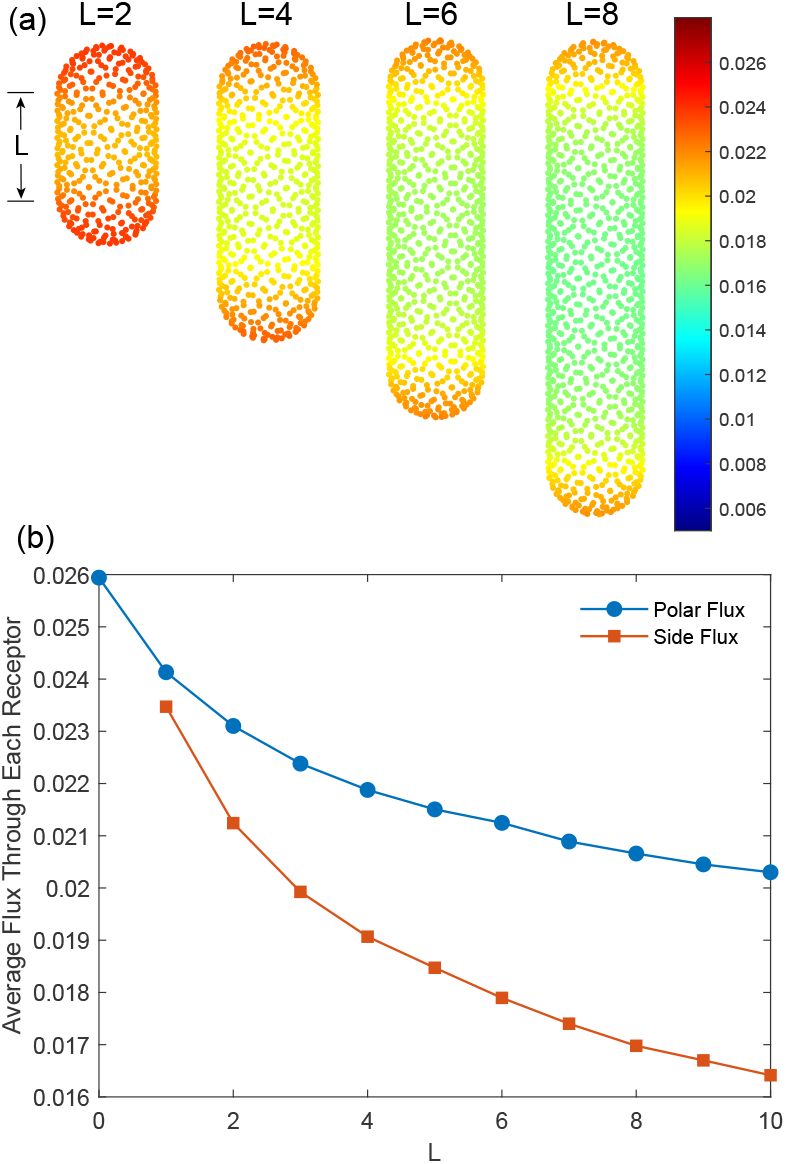
(a) We uniformly sample *N* receptors on a rod-shaped cell, modeled as a cylinder with length *L*, capped with two half-spheres with radius *R* = 1. Here, we show the numerical results of four cells where *N* = 400 for *L* = 2, *N* = 600 for *L* = 4, *N* = 800 for *L* = 6, and *N* = 1000 for *L* = 8. (b) The average fluxes for receptors on the side and polar regions. As the cylinder length increases, the difference between the polar and side flux becomes more significant. In this work, we set the receptor radius *r* = 0.01 unless otherwise mentioned.

For curved rod-shaped cells, an approximation of bacteria such as *Caulobacter crescentus*, we find that the fluxes on the negative-curvature side are lower than those on the other side (Figure 3a,b). We also study spherical cells with an invagination (Figure 3c) and a bud (Figure 3d). Interestingly, for the spherical cell with an invagination, the fluxes are the highest at the edge of the invagination. This observation aligns with the experimental observations that the PTS proteins are localized at the edge of the invagination in defective spherical *E. coli* cells [13]. Our theories unveil the benefit of such a localization pattern. For the spherical cell with a bud, the fluxes are the highest at the tip of the bud and lowest in the neck, connecting the body and bud. Our results suggest that receptors may localize in the bud tip for cells such as budding yeast to enhance the total flux.

**FIG. 3.**
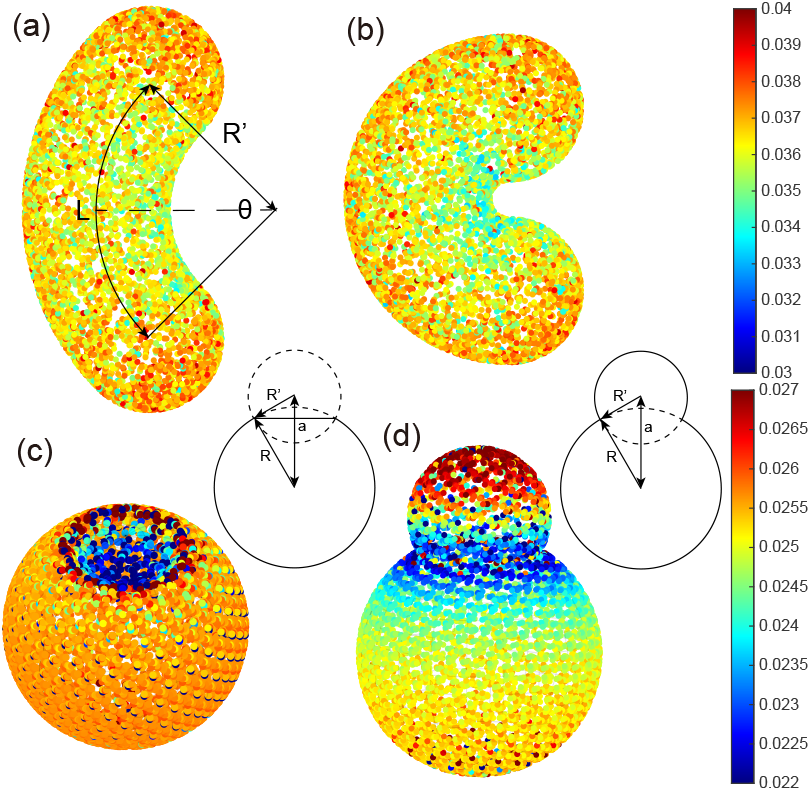
(a-b) We model curved rod-shaped cells as a curved cylinder of contour length *L* = 4 capped with two half spheres with radius *R* = 1. The curved cylinder has a radius of curvature *R*^′^ = *L/θ*, where *θ* = *π/*2 for (a) and *θ* = *π* for (b). For this shape, it is hard to sample receptors uniformly; therefore, we randomly sample *N* = 300 receptors and *M* = 300 auxiliary points with a uniform density, justified by the spherical case (Figure 1b). We accumulate the numerical results of 25 independent samplings. (c) We uniformly sample *N* = 407 receptors on a defective spherical cell. We accumulate the results of 20 independent samplings to show the high fluxes at the invagination edge more clearly. (d) We uniformly sample *N* = 400 receptors on a spherical cell with a bud and accumulate the results of 20 independent samplings. The geometry of the cells <u>i</u>n (c) and (d) is shown on the right of each figure, in which 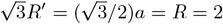.

The spatial dependence of fluxes makes us wonder if we allow the locations of receptors to change from a uniform distribution, is there an optimal distribution that maximizes the total flux? We answer this question in the following section.

## The optimal distribution of receptors on an arbitrarily shaped cell

We next seek the optimal distribution of receptors on an arbitrarily shaped cell that maximizes the total flux. For the optimal distribution, the flux through each receptor must be equal. If not, the cell can always let the receptors move toward the regions with higher fluxes to enhance the total flux. First, we point out a useful connection between the flux through a receptor and the concentration near the receptor: *c*_*i*_ = *J*_*i*_*/*4*rD*. To see this, we let ***x***_*j*_ near ***y***_*i*_ (*r*≪|***x***_*j*_ −***y***_*i*_| ∼O(*l*)) and take the difference between Eq. (4) and Eq. (5), from which we obtain *c*_*i*_ = *J*_*i*_*/*4*rD* + O(1*/l*) ≈ *J*_*i*_*/*4*rD*. Given the fact that *J*_1_ = *J*_2_ = … = *J* for optimal receptor distributions, we have 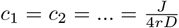 for all receptors. Since the concentrations near all receptors are the same, the concentration over the entire surface should be approximately constant as well, i.e., 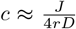.This allows us to simplify Eq. (2) to:

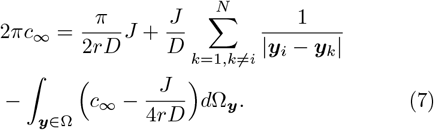

Here, 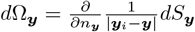,the differential solid angle at ***y*** towards ***y***_*i*_.

Let *σ*(***y***) be the local density of receptors at ***y***. Using 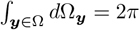,we derive from Eq. (7) that for arbitrary ***y***_*i*_ ∈ Ω:

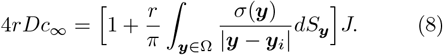

Thus, for ***x*** ∈ Ω:

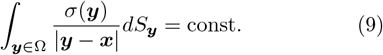

This corresponds to the isopotential condition of ideal conductors if we take *σ*(***y***) as the charge density. Therefore, we predict that the density distribution of receptors maximizing the total flux is the same as the charge density distribution on an ideal conductor of the same shape. To test this prediction, we simulate an evolutionary process on rod-shaped cells and cells with a bud, in which we let the locations of receptors evolve to maximize the total flux. At each step, we let a few receptors change their locations and accept the change if the total flux increases. We confirm that the fluxes are approximately the same across different receptors for the optimal distributions (Figure 4). We then compare the optimal receptor distribution with the charge density distribution on an ideal conductor of the same shape. For both rod-shaped cells and budding cells, the optimal receptor distributions are indeed the same as the charge density distributions on an ideal conductor of the same shape (Figure 4), confirming our prediction.

**FIG. 4.**
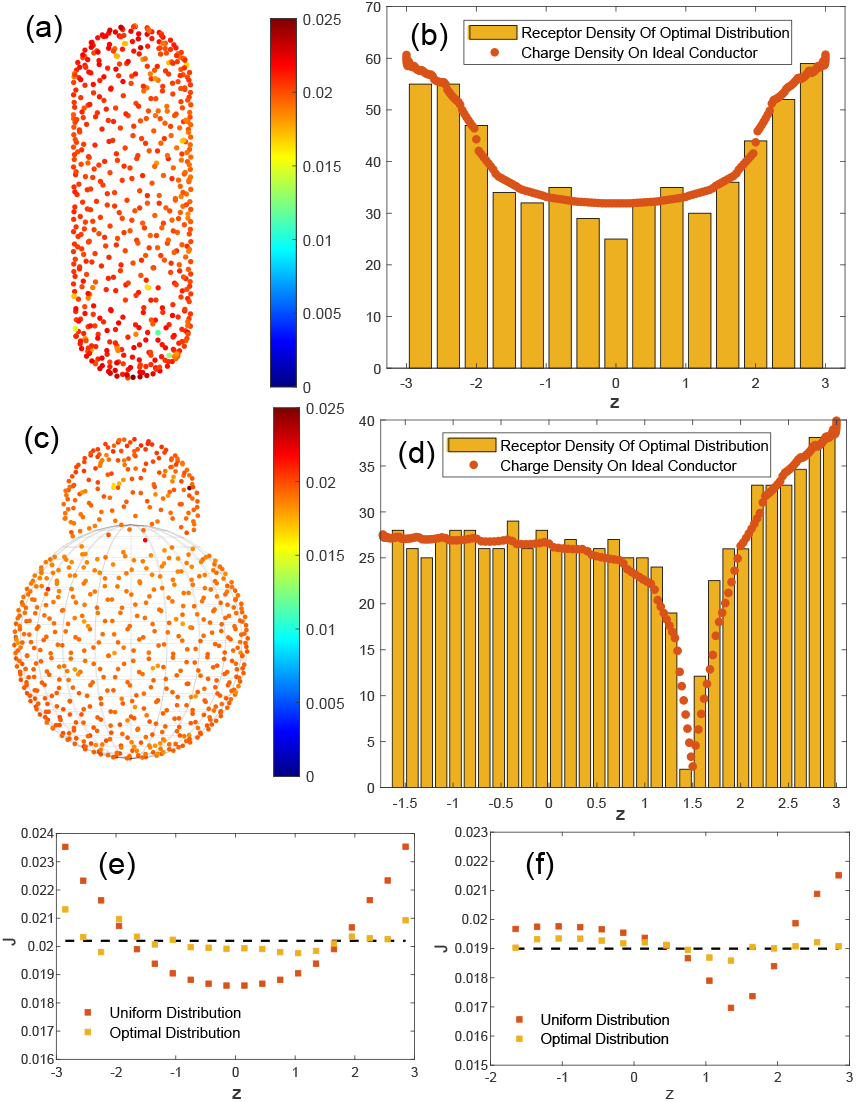
Optimal distributions of receptors on a rod-shaped cell and a cell with a bud. (a) The optimal receptor distribution on a rod-shaped cell with *L* = 4 and *R* = 1. We uniformly distribute 600 receptors initially and let the locations of receptors evolve to maximize the total flux for 5000 time steps. At each time step, we change the positions of 10 randomly chosen receptors and accept the change if the total flux increases. (b) The optimal receptor distribution on the rod-shaped cell agrees with the charge density of an ideal conductor of the same shape. (c) The optimal receptor distribution on a cell of radius √3 and with a bud of radius 1. We uniformly distribute 710 receptors initially and let the locations of receptors evolve to maximize the total flux for 5000 time steps. At each time step, we change the positions of 5 receptors with minimum fluxes and accept the change if the total flux increases. (d) The optimal receptor distribution on the cell surface with a bud agrees with the charge density of an ideal conductor of the same shape. (e-f) The fluxes *J* for receptors with different locations along the vertical direction of the rod-shaped cell (e) and the budding cell (f). For uniformly distributed receptors, the fluxes vary significantly. In contrast, the spatial heterogeneity of fluxes is much smaller for optimally distributed receptors.

## Discussions

In this work, we innovate a new numerical method that converts solving a complex Poisson equation with mixed boundary conditions to a set of linear equations, which allows us to compute the fluxes of all receptors exceedingly fast. Our method not only recovers the classical results of a spherical cell [18] but is also generalizable to arbitrarily shaped cells. Our calculations provide an explanation for several spatial patterns of receptors observed in experiments, including localizations near polar regions [8–12] and invagination [13]. Remarkably, we demonstrate that for an arbitrarily shaped cell, the optimal distribution of receptors on a cell surface is precisely the charge density distribution on an ideal conductor of the same shape, which is a novel prediction as far as we realize.

We provide an intuitive picture underlying the analogy between the optimal receptor distribution and the charge density of an ideal conductor. In the case of optimal receptor distributions, the fluxes through all receptors are the same. Therefore, if we average the fluxes of the receptors inside a patch of size *ξ* such that *l* ≪*ξ* ≪*R*, the direction of the average flux is perpendicular to the cell surface (if the fluxes of the receptors in this patch are unequal, the average flux would be tilted relative to the normal direction of the cell surface). Because the perpendicular nature of the mesoscopic fluxes agrees precisely with the perpendicular property of the electric field relative to the ideal conductor surface, we expect the receptor distribution to be the same as the charge density of an ideal conductor.

We thank Kaifeng Weng for providing us the charge densities of ideal conductors. The research was funded by the National Key Research and Development Program of China (2024YFA0919600) and supported by PekingTsinghua Center for Life Sciences grants.

## Notes

### Competing Interest Statement

The authors have declared no competing interest.

